# Visual featural topography in the human ventral visual pathway

**DOI:** 10.1101/2020.03.27.009498

**Authors:** Shijia Fan, Xiaosha Wang, Xiaoying Wang, Tao Wei, Yanchao Bi

**Author notes:** Corresponding author (YB).

## Abstract

Visual object recognition in humans and nonhuman primates is achieved by the ventral visual pathway (ventral occipital-temporal cortex, VOTC), which shows a well-documented object domain structure. An on-going question has been what type of information is processed in higher-order VOTC that underlies such observations, with recent evidence suggesting effects of certain visual features. Combining computational vision models, fMRI experiment using a parametric-modulation approach, and natural image statistics of common objects, we depicted the neural distribution of a comprehensive set of visual features in VOTC, identifying voxel sensitivities to specific feature sets across geometry/shape, Fourier power, and color. The visual feature combination pattern in VOTC is significantly explained by their relationships to different types of response-action computation (Fight-or-Flight, Navigation, and Manipulation), as derived from behavioral ratings and natural image statistics. These results offer the first comprehensive visual featural map in VOTC and a plausible theoretical explanation as a mapping onto different types of downstream response-action systems.

## Introduction

Ventral occipital-temporal cortex (VOTC), which underlies visual object recognition in humans and nonhuman primates, has a hierarchical architecture, from a retinotopic organization of simple features in the early visual cortex to a domain-based organization (e.g., animate vs. inanimate; faces vs. scenes) in higher-order visual cortex [1–5]. One of the key questions is about the nature of representation in higher level VOTC [6–12]: Exactly what kinds of information about or associated with these various object domains are represented here?

Being the higher-order “visual” cortex, a major candidate representation being assumed is visual features [13]. The effects of two specific types of visual features that associate with the object domains in higher-order VOTC have been very recently demonstrated in humans and nonhuman primates – mid-level shapes and colors. For mid-level shapes, high rectilinearity, especially right angles, is more prevalent in images of scenes and places and activates scene-preferring regions including the parahippocampal place area (PPA) and transverse occipital sulcus more strongly than curved lines in humans [14]. Low rectilinearity, or high curvature, tends to be associated with animate items [15,16], and tends to activate regions close to the face patches in the macaque brain [17]. Different colors have been also shown to associate with objects versus their backgrounds, and with animate versus inanimate objects [18]. Three VOTC patches were identified in the macaque monkeys to be sensitive to color and the more anterior medial patch showed both a yellow/red preference and face/body preference [19]. These studies focus on individual visual features, and it is unknown whether the specific featural effects are driven by other features that correlate with them in various object contexts. Furthermore, to what extent a single feature could explain the observed VOTC domain distribution is controversial [20]: the anatomical overlap between feature effects and domain effects is far from perfect [17], and the domain preferences are still present when visual shape are controlled [8,10]. Our first aim, then, is to depict a comprehensive topographical map of visual features across VOTC, taking into consideration their correlational nature in the context of common objects.

The harder question is, if there were a systematic pattern of various visual feature sensitivity across VOTC, what factor drives this organization. That is, why does a certain region prefer a particular feature or set of features together, or why are various features preferred by the same, or different, brain region in a particular location? Note that “domain preference” in VOTC describes the phenomenon and does not constitute a satisfying explanatory variable for the potential featural effects here, because what constitutes the “domain” information being represented is not explicit. A recent proposal is that the neuronal functional preference of VOTC voxels is constrained, at least partly, by the connectivity pattern with the downstream nonvisual, response-action computations for objects such as Fight-or-Flight, Navigation and Manipulation [12,21–24]. This notion predicts that the visual feature distribution pattern in VOTC is driven by how they may associate with various salient response-action systems in the real world, and will be tested empirically as the second aim of our study.

To depict a comprehensive topographical map of visual features across VOTC, we combined computational vision modelling and parametric modulation analysis on fMRI responses. The parametric modulation approach exploits the natural variation in salience of various visual features across object images (obtained from computational vision modelling) and identifies brain regions responsive to each feature or combination of features by computing the degree of association between brain responses and image feature weights. To test the potential hypothesis about the driving forces of the VOTC visual featural organization, we examined what prototypical visual features are associated with the major types of response-actions to objects (Fight-or-Flight, Navigation and Manipulation), using behavioral rating and natural image statistics in broader image sets, and then examined how well they align with the preferred-visual-feature-sets across various VOTC regions. Finally, the extent to which the documented object domain pattern might reflect the identified visual feature maps was tested using an independent fMRI dataset.

## Results

Twenty visual features covering a broad range of shape, spatial frequency, orientation, and color information were tested, and their weights were extracted for each of 95 object images using computational vision models (see Materials and Methods and **S1 Fig** for details) [14,17,25–29]. fMRI responses for these images were also obtained from 26 participants, and parametric modulation models were used to compute the effects of visual features across VOTC voxels, taking into consideration their inter-correlations (see **Fig 1**). Then an explicit theoretical hypothesis for VOTC computation (visual-feature-for-action response mapping) was tested for the explanatory power for the VOTC visual feature patterns. The relation between featural effects and the domain effects was also examined.

**Fig 1.**
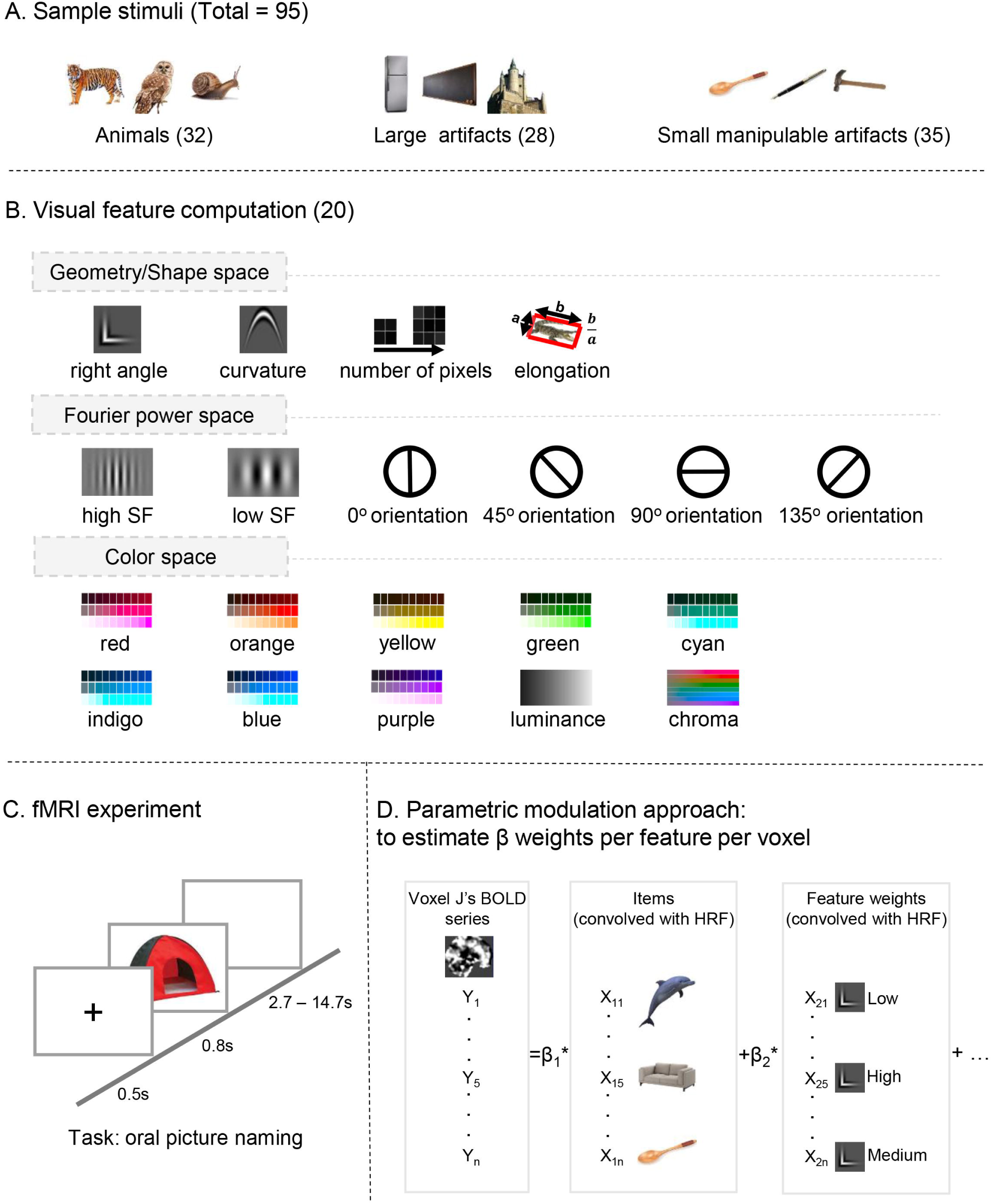
Schematic overview of the methods in main fMRI experiment. (A) Sample stimuli. Images of 95 common objects (32 animate items and 63 inanimate items, including 28 large artifacts and 35 small manipulable artifacts) were used. (B) Visual feature construction from computational vision models. For each picture, computational vision models were used to obtain values of 20 visual features, including geometry/shape (based on modified Gabor filters), Fourier power features (using 2D fast Fourier transform), color (based on Cmmission Internationale de l’Eclairage (CIE) L*C*H space). See **S1 Fig** and Materials and Methods for model construction method details. (C) fMRI experiment. In an event-related fMRI experiment, participants viewed and named these objects. (D) Parametric modulation analysis. Parametric modulation was used to estimate the degree of association between brain responses and visual feature weights across the whole VOTC.

### Computation of visual feature weights in object images

A set of 95 real object images (32 animate items and 63 inanimate artifacts, including 28 large artifacts and 35 small manipulable artifacts) were analyzed using computational vision models to obtain their properties for 20 visual features (see Materials and Methods): in geometry/shape space these features were right angle, curvature, number of pixels and elongation; in Fourier power space high/low spatial frequencies and four orientations (0, 45, 90, 135°); in color space eight hues,luminance, and chroma. The descriptive statistics, including distribution plots for each feature across the whole image set, as well as the mean and standard deviation (SD) by domains, are shown in **S2 Fig**. The correlations (Pearson) among features are shown in **S3 Fig**. As often observed, we found significant differences between animate items and inanimate artifacts (Welch t-test and FDR corrected *q* < .05) across three visual features: right angle (t(63.41) = -3.96, *p* = 1.90 x 10^−4^), elongation (t(68.60) = -3.97, *p* = 1.74 x 10^−4^), and 135° orientation (t(39.85) = 3.12, *p* = 3.33 x 10^−3^). When separating the inanimate objects further into large artifacts and small manipulable artifacts, more features exhibited significant between-domain differences (one-way ANOVA and FDR corrected *q* < .05): right angle (F(2,92) = 6.77, *p* = .002), number of pixels (F(2,92) = 16.37, *p* = 8.27× 10^−7^) and elongation (F(2,92) = 15.47, *p* = 1.61 × 10^−6^) in geometry/shape space; low spatial frequency (F(2,92) = 6.59, *p* = .002), 0° orientation (F(2,92) = 6.21, *p* = .003), 90° orientation (F(2,92) = 5.08, *p* = .008) and 135° orientation (F(2,92) = 8.06, *p* = .001) in Fourier power space; orange (F(2,92) = 5.11, *p* = .008) and yellow (F(2,92) = 5.43, *p* = .006) in color space. The post-hoc comparisons across domain pairs are shown in **S1 Table**. Pairs of highly-correlated visual features (Pearson *r* > .85) were collapsed into one by taking the means (cyan/indigo, *r* = .92, red/purple, *r* = .86). To reduce chances of multicollinearity, low spatial frequency was further excluded from the full parametric modulation model analysis because it had a variance inflation factor (VIF) > 10 [30] (VIF = 48.25; other features’ VIFs are within the range of 1.26 – 5.41). Thus, 17 features were retained for the subsequent parametric modulation analysis, with pairwise correlations within the range of -.56 to .64.

### Visual featural topography map in VOTC

For all fMRI results below, we adopted a threshold of cluster-level FWE corrected *p* < .05 within the VOTC mask [31], with voxel-wise *p* < .001 unless explicitly stated otherwise.

The results of the full model analysis, where the 17 visual feature weights were entered into the parametric modulation model for BOLD activity estimates, are shown in **Fig 2**. In higher-order VOTC, for geometry/shape-space features, right angle modulated responses in bilateral medial fusiform gyrus (medFG) and left lateral occipital temporal cortex (LOTC); number of pixels modulated responses in left medFG. For Fourier-power-space features, high spatial frequency modulated responses in bilateral medFG; 0° orientation modulated responses in right medFG and bilateral LOTC; oblique orientations (45°, 135°) modulated responses in right lateral fusiform gyrus (latFG) and orientation 135° additionally modulated responses in left latFG. For color-space features, red/purple and green modulated broad regions in bilateral FG; red/purple additionally modulated responses in left LOTC; luminance modulated responses in bilateral latFG.

**Fig 2.**
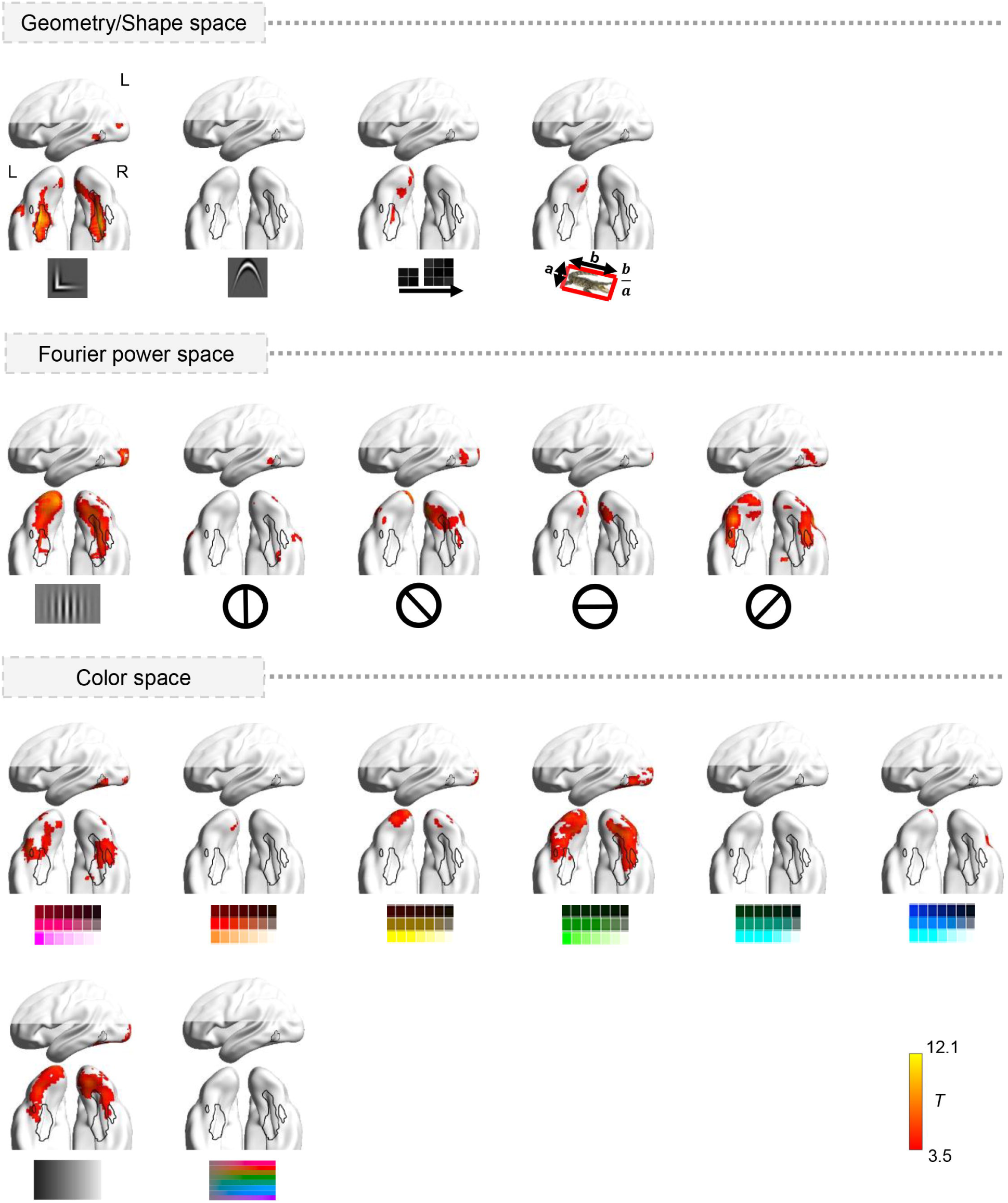
Object visual feature topography in a full-model parametric modulation analysis. All visual feature weights were entered into the parametric modulation model for BOLD activity estimates, yielding an activation map for each visual feature in the VOTC mask. The maps are thresholded at cluster-level FWE corrected *p* < .05 within the VOTC mask, with voxel-wise *p* < .001. The outlines show the object-domain-preferring clusters for animals (bilateral latFG), large artifacts (bilateral PPA), and small manipulable artifacts (left LOTC), localized by contrasting each object domain with the other two domains in the main fMRI experiment.

Independent models, in which each feature was entered into the parametric modulation model separately without considering the correlations among features, were also performed and are shown in **S4 Fig**. Here more commonalities across features can be observed, with most features showing regions largely consistent with those obtained in the full model above with effects covering broader regions in higher-order VOTC. Five features showed differences between the two analyses: The effects of elongation (in left LOTC), 90° orientation (in bilateral medFG), and blue (in bilateral medFG) were significant in the independent model but not in the full model; the effects of 0° orientation (in bilateral LOTC) and luminance (in bilateral latFG) were significant in the full model but not in the independent model. These differences are likely due to their correlations with other features (see **S3 Fig**).

### Factors driving the feature distribution patterns in VOTC voxels

We have described the distributional topography of a comprehensive set of visual features in VOTC, identifying voxels by their sensitivities to specific feature sets. The follow-up question is why visual features distribute in this way. Why does a certain region prefer a particular feature set together (e.g., right angle, high spatial frequency, and 0° orientation in medFG), or why does a particular feature set is preferred by the same brain region in a particular location? We test an explicit notion that the neuronal functional preference of VOTC voxels to certain visual features is constrained, at least partly, by the connectivity pattern with the downstream computations [12,21–24]. That is, the pattern of visual features associated with various types of response-actions in real world constrains the visual feature distribution pattern in VOTC. We first obtained the type of visual feature clustering patterns associated with the nonvisual response-action properties by behavioral ratings and computations of natural images (procedure in **Fig 3A**; Results in the radial-maps in **Fig 3B**, left column). A binary-labeled “animacy domain” model was also tested as a reference (**S5 Fig**, left column). We then tested whether the response- and animacy-domain categorization-associated visual featural patterns indeed align with the visual feature topography of VOTC (**Fig 3B** and **S5 Fig**, brain figures).

**Fig 3.**
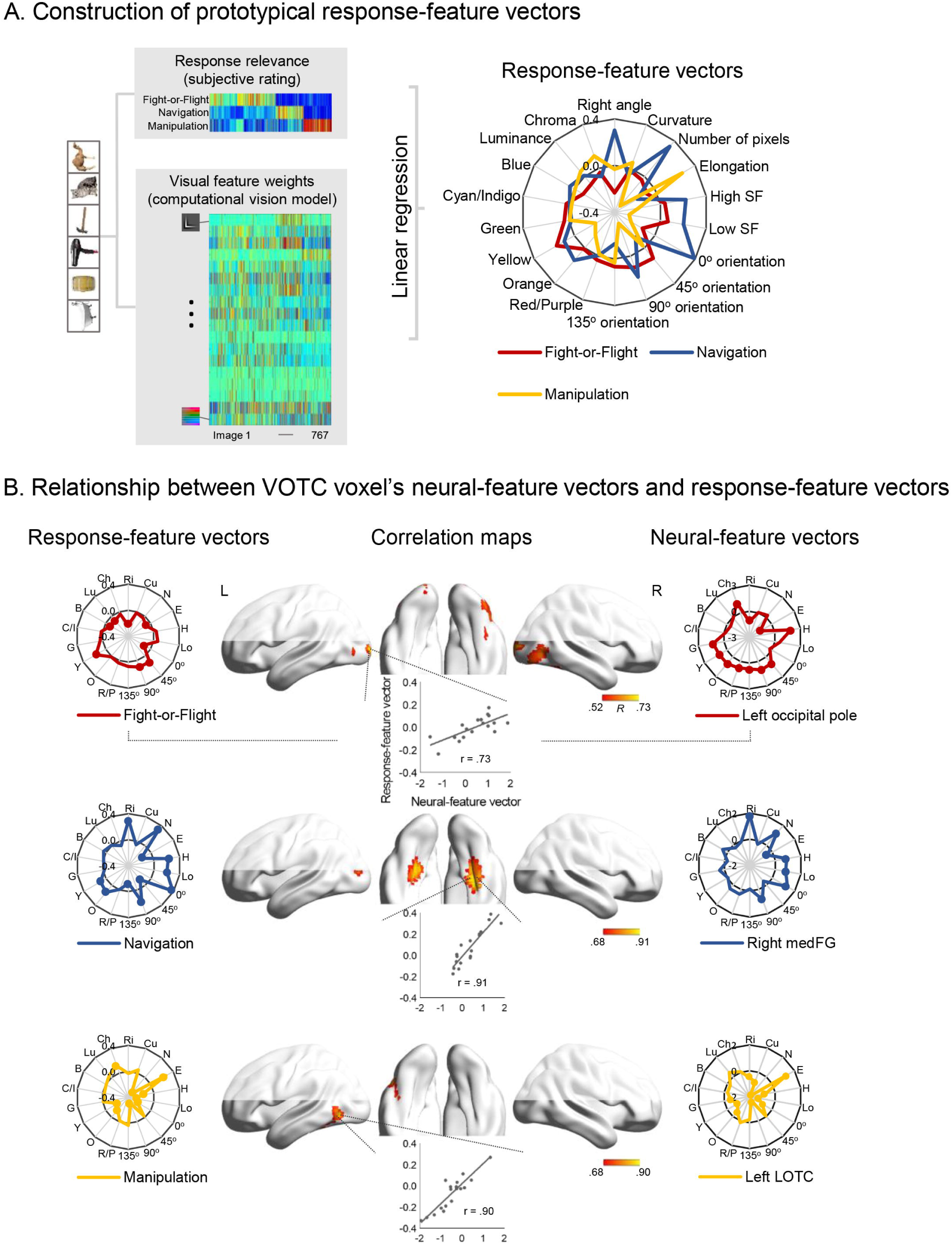
Relationship between response-driven-model and VOTC’s visual featural topography. (A) Illustration of the construction scheme of prototypical visual-feature vectors for the three response-action systems. In an image set of 767 images, visual feature weights for each image were obtained using computational vision models. We examined 3 theorized response systems (Fight-or-Flight, Navigation and Manipulation) by asking 24 participants to rate how strongly each object associates with each of the three response-action systems. Linear regressions were conducted between each response vector and each visual feature weight, resulting in 3 response-feature beta vectors. In (B), the left column shows the “prototypical” visual feature vectors associated with each response-action system (Fight-or-Flight, Navigation and Manipulation). Dots indicate that beta values were significant at FDR corrected *q* < .05 for 54 comparisons. The middle column shows the Pearson correlation maps between each of these “prototypical” response-driven-visual vectors and VOTC voxels’ neural -feature vector obtained from the fMRI parametric modulation analyses. The correlation maps are thresholded at cluster-level FWE corrected *p* < .05, voxel-wise *p* < .001 for Navigation-driven and Manipulation-driven vector, and voxel-wise *p* < .01, cluster size > 10 for Fight/Flight-driven vector. Scatter plots show the correlations for the peak voxels. The right column presents the peak voxel’s neural-feature vectors.

#### Relationship between observed VOTC voxel feature vectors and response-feature vectors

We obtained prototypical visual feature vectors associated with hypothesized response-action systems (i.e., response-feature vector) based on subjective rating and natural image statistics to approximate the feature-response association profile in the real world. For this purpose, we used a broader image set containing 767 images, which included the 95 images from the current fMRI experiment and 672 images (isolated objects with clear domain membership on white background) selected from three previous studies [32–34]. Three types of response-actions were considered:

Fight-or-Flight, Navigation and Manipulation [12,21,23]. We first asked an independent group of participants to rate each object image on its relevance to each response (see Materials and Methods for details). Linear regression was conducted between the rated value for each response-action system and each of the 18 visual feature weights; the resulting beta values were used as the prototypical response-feature vector for each response-action system in natural object images. The results (FDR corrected *q* < .05) are shown in **Fig 3B**, left column (see **S2 Table** for beta and *p*-values).

We then tested whether and in what manner the VOTC voxels’ visual feature sensitivity profiles align with these response-associated-feature vectors. For each VOTC voxel we correlated its neural-feature vector with each of the three prototypical response-feature vectors, resulting in 3 VOTC correlation maps (**Fig 3B**, brain figures). Note that the independent parametric modulation model results were used (with 18 visual features), because the beta weights were more transparently interpretable and the results were also largely consistent with those of the full model (17 features, with “low spatial frequency” excluded due to high VIF). The navigation-response-feature vector was significantly correlated with the neural-feature vector in clusters located in bilateral medFG, left middle occipital gyrus and right lingual gyrus. The manipulation-response-feature vector was significantly correlated with the neural-feature vector of VOTC voxels in clusters located in left LOTC and left lingual gyrus. The fight/flight-response-feature vector showed no significant correlations at the standard threshold. When we lowered the threshold (voxel-wise *p* < .01, cluster size > 10), it correlated with the neural-feature vector of VOTC voxels in bilateral lateral occipital cortex, right latFG and bilateral occipital pole.

#### Relationship between observed VOTC voxel feature vectors and animacy-domain-feature vectors

Although the specific computation goals served by the “domain” organization (i.e., driving factor) are not articulated, we nonetheless generated prototypical visual feature vectors associated with the widely proposed categorization (i.e., domain-feature vectors) based on natural image statistics to approximate the feature-domain association profile in the real world. Binary domain labels were constructed (for animacy: animate items = 1, inanimate items = 0). Logistic regression was conducted between each binary domain label and each of visual feature weights across 767 images mentioned above, and the resulting beta values were taken as a prototypical domain-feature vector for each domain. This vector reflects the visual feature patterns that best distinguish animate and inanimate items in natural object images. The results (FDR corrected *q* < .05) are shown in **S5 Fig**, left column; see **S2 Table** for beta and *p*-values). We then tested whether the VOTC voxels’ visual feature sensitivity profiles reflect these animacy-domain-feature vectors. For each VOTC voxel, we correlated its neural-feature vector (obtained by the independent parametric modulation model; **S4 Fig**) and each of the two prototypical domain-feature vectors, resulting in a correlation map for each domain (**S5 Fig**, brain figures). The animate-domain-feature vector was significantly correlated with the neural-feature vector of VOTC voxels in one cluster located in right lateral occipital cortex. The inanimate-domain-feature vector was significantly correlated with the neural-feature vector of VOTC voxels in three clusters located in bilateral medFG and left LOTC. These results suggest that VOTC voxels’ feature-sensitivity patterns are associated with the natural image statistics of two major object domains.

#### Comparison between response-driven and animacy-domain-driven hypotheses

We directly compared the explanatory power of these two types of feature vectors to see if, by being more specific, the response-driven model captures finer properties of the VOTC visual feature topography. To do this, we first generated a response-driven maximum R map by selecting the highest R value for each voxel out of the three response-driven R maps in **Fig 3B**, and generated the animacy-domain-driven maximum R map in the same way using the two maps in **S5 Fig**. Then the two max R maps were Fisher-z transformed and compared by paired t-test. Results showed that the “response-driven” map was significantly higher than the “animacy-domain-driven” map (global mean Rs ± SD: 0.57 ± 0.27 vs. 0.39 ± 0.23, *t* (3914) = 34.87, *p* = 2.87 x 10^−232^).

Inanimate artifacts have been further divided into large artifacts and small manipulable artifacts in recent studies which showed a tripartite structure of large artifacts, animals, and small manipulable artifacts in VOTC, spanning from the medial fusiform/parahippocampal gyrus to the LOTC [4,31,35]. We also tested whether response-driven model has greater explanatory power than the tripartite-domain-driven model because they are highly correspondent (Fight-or-Flight responses with animals; Navigation with large artifacts; Manipulation with small manipulable artifacts). Similar procedure as the previous analysis was repeated and results showed that the “response-driven” maximum R map was still significantly higher than the “tripartite-domain-driven “maximum R map (global mean Rs ± SD: 0.57 ± 0.27 vs. 0.54 ± 0.28, *t* (3914) =12.21, *p* = 1.07 x 10^−33^; See **S6 Fig** for results of prototypical tripartite-domain-feature vectors and correlation maps).

#### Association between visual featural effects and the domain effects in VOTC

Having established the visual featural topography in VOTC and tested the driving variables for such distributions, here we assess to what extent the well-established object-domain observations (i.e., animacy and size) can be accounted for by the underlying featural representations.

A multiple linear regression model was constructed to predict a voxel’s selectivity strength for object domains (obtained from an independent fMRI dataset) using its visual feature sensitivity patterns, across all VOTC voxels. That is, the 17-feature sensitivity maps in VOTC from the full parametric modulation model were taken as the independent variables. The dependent variable was the VOTC animacy-domain-selectivity strength map obtained from an independent dataset (contrasting animate items with inanimate items; see details in [35,36]). The results (**Fig 4A)** showed high explanatory power of the linear regression model: adjusted-R^2^ = .815. Using the animacy-domain-selectivity strength map computed from the main fMRI experiment data with the identical contrast (i.e., within-subject analysis) yielded an adjusted-R^2^ of .959.

**Fig 4.**
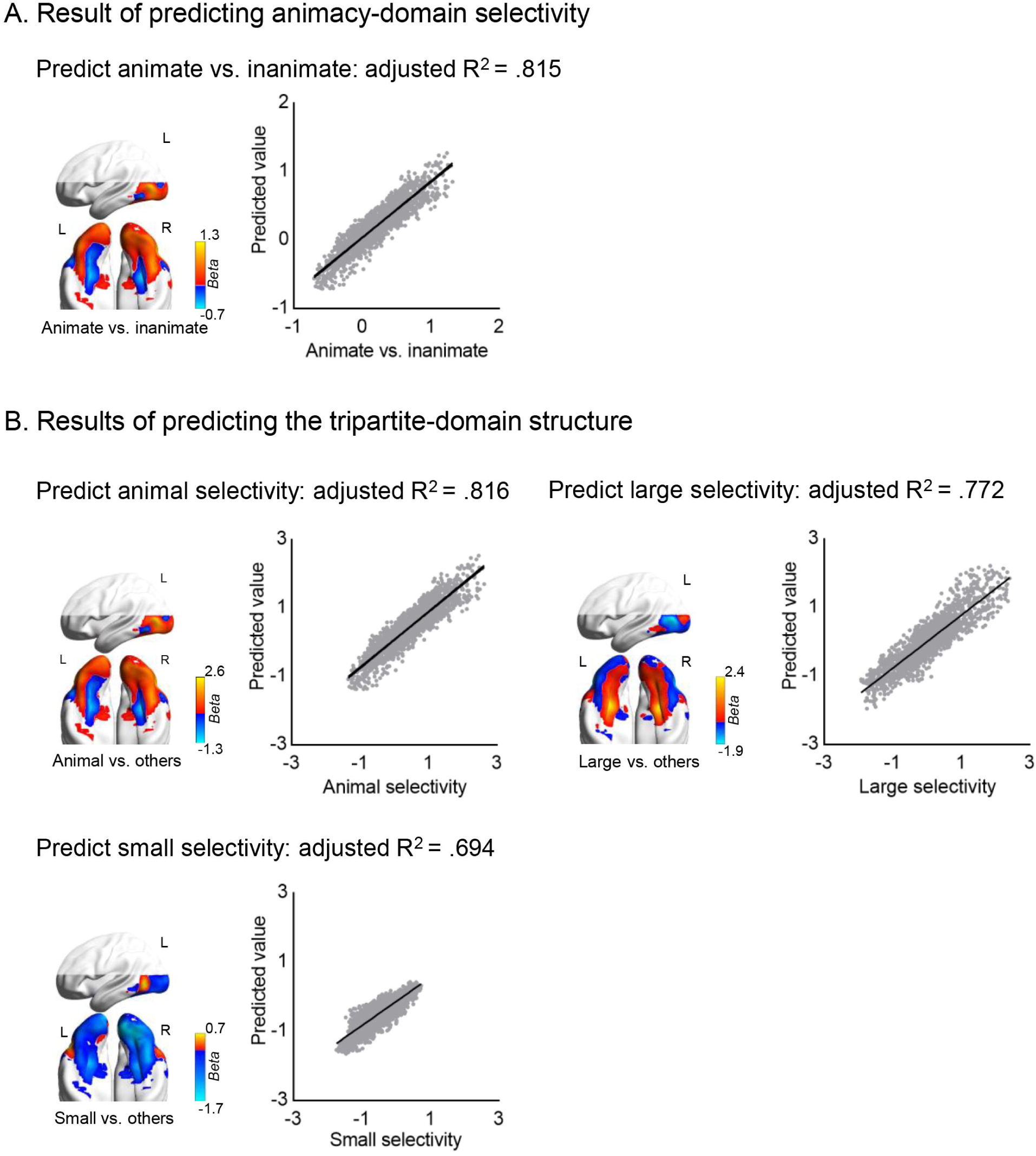
The association between visual feature topography and object domain effects. (A) Result of using VOTC’s visual feature vectors to predict animacy-domain selectivity: A multiple linear regression model was constructed to predict domain selectivity strength for animate/inanimate domains, using 17 visual features’ beta values as predictors, across all VOTC voxels. The brain map is the unthresholded animate vs. inanimate activation map, showing the group-averaged selectivity strength (beta values of animate items – inanimate items for all VOTC voxels). The scatter plot shows correlation between predicted animacy-domain-selectivity strength using VOTC visual-feature maps and the observed domain-selectivity strength across all VOTC voxels. (B) Results of using visual feature vectors to predict the tripartite-domain structure: Three multiple linear regression models were constructed to predict domain-selectivity strength for animals, large artifacts or small manipulable artifacts, using 17 visual features’ beta values as predictors, across all VOTC voxels. The brain maps are the unthresholded activation maps for animals, large artifacts or small manipulable artifacts, showing the group-averaged selectivity strength (beta values of one domain – those of other two) for all VOTC voxels. The scatter plots show correlations between predicted domain-selectivity strength using VOTC visual-feature maps and the observed domain-selectivity strength across all VOTC voxels.

For predicting the tripartite-structure (animals, large artifacts, small manipulable artifacts), the dependent variable was obtained by contrasting beta values of each domain with the mean of the other two. Again, using the independent dataset, a voxel’s visual feature vector can highly significantly predict its selectivity strength (**Fig 4B**) for animals (adjusted-R^2^ = .816), for large artifacts (adjusted-R^2^ = .772), and for small manipulable artifacts (adjusted-R^2^ = .694). The results were higher using data from the same main fMRI experiment (for animals, adjusted-R^2^ = .957; for large artifacts selectivity, adjusted-R^2^ = .946; for small manipulable artifacts selectivity, adjusted-R^2^ = .973).

## Discussion

Combining computational vision models, a parametric modulation analysis of fMRI data, and natural image statistics, we depicted the distributional topography of a comprehensive set of visual features (geometry/shape, Fourier power, and color) in VOTC, identifying voxels’ sensitivities to specific feature sets. We demonstrated that the relationship with salient response actions in real world can explain why visual features are distributed this way in VOTC.

By contrast to recent studies that focused on one or two specific visual features or unarticulated DNN-derived hidden spaces [13,37], our approach tested a much more comprehensive set of visual features and correlations among them, and showed highly significant explanatory power for the well-documented object domain structure. Previous studies have shown associations between certain visual feature and domain preferences: A preference for rectilinearity, high spatial frequency, and cardinal orientation features was observed in regions preferring scenes/large objects [14,26,38,39] and a preference for high curvature, low spatial frequency, and red/yellow hues was observed in regions preferring faces [17,19,26,40]. However, these visual features do not explain the domain observations satisfactorily, in terms of the selectivity strengths [20], the anatomical overlap [17], and the domain preference effects that are still present when visual shape are controlled [10]. Here, by incorporating the combinational effects of multiple visual features together, we showed remarkably high explanatory power of visual features to domain-preference: Voxels’ visual-feature-preference vectors accounted for over 80% of the variance in VOTC’s selectivity for the animate/inanimate domain selectivity, and over 69% of the variance in selectivity for animals, large artifacts or small manipulable artifacts. Our results not only provide a computational model that theoretically may predict the VOTC neural activity pattern for objects based on its visual featural properties, including those along fuzzy domain boundaries, but also offer positive evidence for a plausible, specific representation theory of VOTC that can explain the domain-appearing observations: What VOTC represents (at least) is visual features.

Why does VOTC have this specific type of visual featural topography then? We provided evidence that is consistent with the recent proposal that the neuronal functionality of VOTC voxels is constrained, at least partly, by the association pattern with the downstream nonvisual, action computations such as Fight-or-Flight, Navigation and Manipulation [12,21–24]. Prototypical visual feature sets that associate with the three types of response actions, obtained through rating and natural image statistics, indeed align with those preferred visual feature combination patterns in different patches of VOTC: fight/flight-response vector (3 highest loadings in yellow, 45° orientation, 90° orientation; 3 lowest loadings in right angle, 0° orientation, bluish) was associated with right latFG, bilateral lateral occipital cortex and bilateral occipital pole; navigation-response vector (3 highest loadings in 0° orientation, number of pixels, right angle; 3 lowest loadings in elongation, 135° orientation, 45° orientation) was associated with bilateral medFG, left middle occipital gyrus and right lingual gyrus; and manipulation-response vector (3 highest loadings in elongation, chroma, luminance; 3 lowest loadings in number of pixels, 90° orientation, low spatial frequency) with left LOTC and left lingual gyrus. That is, how visual featural sensitivity is organized in VOTC neurons can be explained by how visual features map with down-stream action responses. Those visual features (combinations) that tend to indicate and associate with a certain action response (e.g., manipulation) are being preferentially processed and represented, together, in regions that are optimally connected with the corresponding action systems [12,21–24].

It should be emphasized that we are interpreting our results as showing that the representation here in VOTC is visual features, organized in a way that allows them to optimize mapping with (i.e., driven by) the response-action programs, and not the action programs themselves. This is in line with the vast literature that VOTC is important for visual processing and that damaging dorsal regions, and not VOTC, leads to action deficits. Also worth noting is that these response models are clearly associated with the object domains that have been used to label the VOTC selectivity [2,4]: fight/fight responses with animals; navigation responses with large objects; manipulation responses with small objects. We are treating this “domain” structure as an observation to be explained instead of an explanatory theory because it is descriptive, vaguely defined, and does not offer hypothesis about exactly what information is represented here. The visual-feature-driven-by-action-mapping account not only explains this observation, but also makes predictions that are consistent with a series of results comparing the feature vs. domain effects in the literature: Objects that do not have prototypical shapes of a domain (e.g., a cup shapes like a cow) are processed by VOTC more similarly to items sharing its surface shape (e.g., animal in this case) and not to those in the same domain (regular cups) [41]; the animate-preferring areas are modulated by how “typical” (human-like) animals are [42]; features without domain contexts may still be able to produce effects [14,17,38]. Our supplementary analyses and feature-validation fMRI experiment analyses provided further support to this last point (**S1 Text; S7 and S8 Figs**): The featural effects were largely present when regressing out domain structure; The effect of right angle in bilateral medFG (aligning with the PPA) was present when the features are shown in isolation without object contexts and/or other features, and even during presentation of objects from non-preferring domains (i.e., when objects are small manipulable artifacts or animals). Interestingly, the effects of other features such as hue and orientation were only observed when presented within objects and disappeared when shown in isolation, indicating that they are processed in combination with other visual features and/or object contexts in VOTC [43].

There are two caveats to consider. One is that the visual features we tested are based on knowledge and algorithms from computational vision practice. There is always a possibility that other relevant types of visual features were missed, and that the algorithm choice was not optimal. For instance, the current curvature computation considers 5 arbitrarily-selected concavity features, and its effects on VOTC based on this computation were not significant yet were visible when using a direct contrast (top 25% amount of curvature – top 25% amount of right angle, **S9 Fig**), more in line with studies using subjective curvature ratings, which may reflect a composite index of various types of curvatures [13]. However, our result that the feature combination model highly significantly predicts the domain-preference strength in VOTC voxels indicate the power of the included features. Second, we only examined the major common objects domains, not testing other classical domains for VOTC: scenes and faces. The current framework makes the same predictions about preferences for these two types of images, which remain to be empirically tested.

To conclude, we found that there are systematic patterns of various visual feature sensitivity across VOTC, offering a comprehensive visual feature topography map. Such visual feature topography can be explained by how features map onto different types of response actions. The object-domain-related observations can be largely explained by voxel sensitivity patterns to the visual features. These findings led us to propose a visual-feature based representation in VOTC, which is driven by their association with nonvisual-action computations (stored elsewhere).

## Materials and Methods

### Participants

Twenty-nine participants (age range 19-32 years; 21 females) participated in the main fMRI experiment. An independent group of 19 participants (age range 19-29 years; 11 females) participated in a feature-validation fMRI experiment. An independent group of 24 participants (age range 18-29 years, 15 females) participated in the rating study. All participants had no history of neurological or psychiatric impairment, had normal or corrected-to-normal vision, were native Chinese speakers, and provided written informed consent. The main fMRI experiment and rating studies were approved by the Institutional Review Board of the State Key Laboratory of Cognitive Neuroscience and Learning, Beijing Normal University (ICBIR_A_0040_008), adhering to the Declaration of Helsinki for research involving human subjects. The feature-validation fMRI experiment was approved by the Institutional Review Board of Department of Psychology, Peking University (#2015-05-04), adhering to the Declaration of Helsinki for research involving human subjects.

### fMRI stimuli

Stimuli in the main fMRI experiment consisted of 95 colorful real-world objects centered on a white background, including animals (32 images) and inanimate manmade artifacts (28 large artifacts, 35 small manipulable artifacts). Images of animals included mammals, birds, reptiles and insects. Images of large artifacts included buildings, furniture, appliances, communal facilities and large transport. Images of small manipulable artifacts included common household tools, kitchen utensils, stationery and accessories. These images were obtained from the Internet and resized to 400 x 400 pixels (10.55° x 10.55° of visual angle).

We also collected a validation dataset that included domain-functional localizer runs and runs of visual features that were presented out of the object contexts (see details in **S1 Text**).

### Computation of visual feature weights in object images

The weights of 20 visual features covering a broad range of shape, spatial frequency, orientation, and color properties were extracted using computational vision models for each of 95 object images (see **S1 Fig** for schematic modelling steps).

#### Geometry/shape space

We examined four geometry/shape features: number of pixels, right angle, curvature, and elongation. For number of pixels, a binary object mask (defined as pixels with grayscale values lower than 240) was created and each pixel in the mask was counted. Overall right angle and curvature information was measured largely following previous approaches with some modification [14,17,29]. Specifically, for right angle, 64 right-angle Gabor filters (using an absolute function [14]) were constructed using 4 spatial scales (1/5, 1/9, 1/15, and 1/27 cycles per pixel) and 16 orientations (22.5°-360° in 22.5° steps). Images were converted to grayscale and edge maps were constructed using Canny edge detection at a threshold of 0.1 [44]. Each edge map was convolved with 64 Gabor filters of different spatial scales and orientations. This produced 64 Gabor coefficient images, which were then normalized by dividing by the mean magnitude of each Gabor filter. For each spatial scale, the largest magnitude across the 16 coefficient images of different orientations was extracted for each pixel to obtain a peak Gabor coefficient image, which was then averaged across all pixels of each image and z-scored across the image set. The resulting Gabor coefficient values for each image were finally averaged across 4 spatial scales and z-scored to provide a single value for each image to represent the amount of right-angle information in that image. For curvature, the same procedure was used using the bank of 320 curved Gabor filters (using a square root function [45], composed of 4 spatial scales, 16 orientations, and 5 levels of curvature (π/256, π/128, π/64, π/32, π/16)), to generate a single value for the amount of overall curvature information for each image. Elongation was measured as the aspect ratio of the rectangle that encloses the object parallel to the object’s longest axis.

#### Fourier power space

Images were converted to grayscale and submitted to a 2D fast Fourier transform (built-in MATLAB function fft2). The high/low spatial frequency and 4 orientations (0, 45, 90 and 135°) were measured based on previous approaches [26,28,38] to parameterize energy variation in Fourier power space. The overall energy of high and low spatial frequency was calculated by averaging the energy of the high (>5 cycles/degree) and low (<1 cycles/degree) band for each image [26]. For orientations, we selected four directions which centered on vertical (0°), left oblique (45°), horizontal (90°), and right oblique (135°) with a bandwidth of 20° [28]. For each orientation range, the energy across spatial frequencies was averaged.

#### Color space

Three main perceptual dimensions of color—luminance, chroma, and hue—were quantified using (CIE) L*C*H space following previous studies [18,25]. Pixel colors in each image were converted from RGB space into (CIE) L*C*H space using the MATLAB “colorspace” package [46]. The white point for the transformation of the image colors was set to D65 (the standard illumination for noon daylight). The luminance and chroma of each image were calculated by averaging these values across pixels within the object. The hue layer was divided into 8 bins with equal width, which started from the 338°-23° bin, in 45° steps, and roughly corresponded to red, orange, yellow, green, cyan, indigo, blue, and purple [25]. The number of pixels in each bin was then counted as the hue-specific measures; pixels that did not belong to objects or were ambiguous (defined as luminance or chroma values less than 10) were excluded.

#### Distribution of visual features across and within object domains

Distribution plots of z-transformed visual feature weights across 95 images were plotted and the mean and standard deviation (SD) by domains were calculated. Welch’s t-test (for unequal variances, conducted using MATLAB) was used to test whether there were significant differences between animate and inanimate objects for each feature. When separating the inanimate objects further into large artifacts and small manipulable artifacts, one-way ANOVA (conducted using SPSS Statistics Software version 26 (IBM)) was used to test whether there were significant differences among three domains for each feature, followed by Tukey’s HSD post hoc comparison. Multiple comparisons across features were corrected using FDR.

### fMRI experiment design

Participants were asked to name the 95 object images overtly and as quickly and accurately as possible. There were 6 runs, each lasting for 528 s. Each image (0.5 s fixation followed by 0.8 s image) was presented once per run. Inter-trial intervals ranged from 2.7-14.7 s. The order of stimuli and length of ITI were optimized using optseq2 (http://surfer.nmr.mgh.harvard.edu/optseq/). The order of items was randomized across runs. Each run started and ended with 10 s of blank screen.

### MRI acquisition and data preprocessing

The main fMRI experiment was conducted at the Beijing Normal University Neuroimaging Center using a 3T Siemens Trio Tim scanner. Functional data were collected using an EPI sequence (33 axial slices, TR = 2000 ms, TE = 30 ms, flip angle = 90°, matrix size = 64 × 64, voxel size = 3 × 3 × 3.5 mm^3^ with gap of 0.7 mm). T1-weighted anatomical images were acquired using a 3D MPRAGE sequence: 144 slices, TR = 2530 ms, TE = 3.39 ms, flip angle = 7°, matrix size = 256 × 256, voxel size = 1.33 × 1 × 1.33 mm^3^.

Functional images were preprocessed and analyzed using Statistical Parametric Mapping (SPM12, http://www.fil.ion.ucl.ac.uk/spm), Statistical Non-parametric Permutation Testing Mapping (SnPM13, http://warwick.ac.uk/snpm), and Data Processing & Analysis of Brain Imaging (DPABI) [47]. The first 5 volumes in each run of the main fMRI experiment and feature-validation experiment were discarded. Image preprocessing included slice time correction, head motion correction, and normalization to the Montreal Neurological Institute (MNI) space using unified segmentation (resampling voxel size = 3 × 3 × 3 mm in the main fMRI experiment; 2 × 2 × 2 mm in the feature-validation experiment), and spatial smoothing with a Gaussian kernel of 6 mm full width at half maximum. Three participants in the main fMRI experiment were excluded from analyses due to excessive head motion (>3 mm maximum translation or 3° rotation).

Statistical analyses were carried out within a bilateral VOTC mask (containing 3915 voxels for 3-mm voxel size) constructed in a previous study [31], which was defined as brain regions activated by the contrast of all objects versus fixation in an object picture perception task in the ventral occipitotemporal cortex. Activation maps for parametric modulation and contrasts between conditions (see below for details) were first created in individual participants and then submitted to group-level random-effects analyses using SnPM13. No variance smoothing was used and 5,000 permutations were performed. A conventional cluster extent-based inference threshold (voxel level at *p* < .001; cluster-level FWE corrected *p* < .05 within VOTC mask) was adopted unless stated explicitly otherwise.

### Visual featural topography in VOTC

To identify brain regions associated with each feature, parametric modulation was employed to investigate the correlations between activity levels and feature weights across the 95 stimulus images in the main fMRI experiment. For the full model that considers the correlations among multiple features, the variance inflation factor (VIF) for each feature was calculated using SPSS Statistics Software version 26 (IBM) and features with VIF above 10 were excluded from analysis to reduce multicollinearity [30]. Then the preprocessed functional images of each participant were entered into a General Linear Model (GLM), which included the onsets of items as one regressor, all features’ weights for each image in the parametric modulation module, and 6 head motion regressors for each run. A high-pass filter cutoff was set at 128 s. Contrast images for each feature versus baseline were then calculated and submitted for random-effects analyses. Because there is no *a priori* expectation that any brain region should become “less” active as the processing demands for a given feature increase, making the interpretation of negative correlations speculative, only positive modulations were reported. To obtain raw feature maps without considering correlations among features, we also conducted parametric modulation analyses for each feature by including one feature at a time in the GLM.

To have reference to landmarks showing well-documented object domain preferences, in the result visualization (**Fig 2**) we marked the object-domain-preferring clusters for animals (bilateral latFG), large artifacts (bilateral PPA), and small manipulable artifacts (left LOTC). A GLM including animals, large artifacts, small manipulable artifacts and 6 head motion regressors for each run was constructed. Contrast images of each object domain with the other two domains were calculated in individual level and submitted to SnPM13 for random-effects analyses. The obtained group-level activation maps were thresholded at cluster-level FWE-corrected *p* < .05 within the VOTC mask with voxel-wise *p* < .0001 for animals and large artifacts, and voxel-wise *p* < .01 for small manipulable artifacts. The details of the identified regions were as follows: for animal > others, the bilateral latFG, 51 voxels; for large artifacts > others, the bilateral PPA, 464 voxels; and for small manipulable artifacts > others, the left LOTC, 93 voxels.

### Factors driving the visual feature distribution patterns in VOTC voxels

After establishing the visual featural topography of VOTC, here we try to understand why visual feature sensitivity is distributed across VOTC voxels in the observed way. To test the feasibility of visual-feature-to-map-with-response-action notion, we first examined what type of visual feature clustering patterns associate with the nonvisual response-action properties by behavioral ratings and computations of natural images. A binary-labeled “domain” model was also tested as a reference. We then tested whether the response-action- and binary domain categorization-associated visual feature combination patterns indeed align with the visual feature organization of VOTC.

#### Prototypical visual-feature vectors for response-actions and domains

To gain an unbiased understanding of feature distribution among objects, we built a larger object image dataset containing 672 images from three previous image sets [32–34] and the 95 images from our main fMRI experiment. We used these image sets because they had isolated objects presented on a white background. One object image was the same in our current experiment and in Downing et al. 2006 and thus only one was included. There were 419 animals (mammals, marine creatures, birds, insects, fish, reptiles) and 348 inanimate manmade artifacts (168 large artifacts and 180 small manipulable artifacts, including buildings, furniture, appliances, communal facilities, large transportation, common household tools, kitchen utensils, accessories). All images were resized to 256 × 256 pixels with 72 DPI using Adobe Photoshop CS6. For each image, the feature weights were measured using computational vision models, as described above for main fMRI experiment stimuli. For response-driven prototypical visual-feature vectors, we examined three theorized response-action systems: Fight-or-Flight, Navigation and Manipulation [12,21,23]. The relevance of the 767 images (set described above) to each response-action system was rated by an independent group of participants (N = 24, age range 18-29 years, 15 females) on a 1 – 5 scale. For Fight-or-Flight, the participants were asked to rate “to what extent the object depicted in image would make you to show a stress-related response, e.g., run away, attack, or freeze.” For Navigation, the participants were asked to rate “to what extent the object depicted in image could offer spatial information to help you explore the environment.” For Manipulation, the participants were asked to rate “to what extent the object depicted in image can be grasped easily and used with one hand.” The ratings were averaged across participants to get one relevance index for each image to each response. Then pairwise linear regression analyses were conducted between each response-action relevance type and each visual feature, resulting in 3 response-feature beta vectors.

For domain-driven prototypical visual-feature vectors, we constructed corresponding domain labels (e.g., for animacy domain model: animates = 1, inanimates = 0; for animal-large artifact-small artifact tripartite domain model: animals = 1, others = 0) and performed pairwise logistic regression analyses between each domain label and each of visual features, resulting in domain-feature beta vectors.

#### Effects of response-driven and domain-driven models for VOTC featural topography

We examined whether the VOTC voxels’ visual feature profiles could be driven by response- and/or domain-feature vectors based on natural image statistics. The VOTC voxels’ visual feature profile was generated by extracting group-averaged parameter estimates of each feature in independent models, which resulted in a neural-feature beta vector for each voxel. The independent model results were used here because the beta values were more transparently interpretable and the results of the independent models are largely consistent with those of the full model. Then the neural-visual feature vectors were correlated with each of three response-feature vectors or the object-domain-feature vectors using Pearson correlation, resulting in three R maps for response-driven and tripartite-domain-driven models, and two R maps for animacy-domain-driven model. The significance of the R maps above zero (one-tailed) was thresholded at cluster-level FWE-corrected *p* < .05 within the VOTC mask with voxel-wise *p* < .001. To compare the explanatory powers between response-driven and domain-driven hypotheses, the three R maps derived from response-feature vectors were collapsed by extracting the highest R value in each voxel in VOTC and the two (or three) R maps derived from domain-feature vectors were also collapsed using the same method. The resulting two maximum R maps were then Fisher-Z transformed and compared using paired t-test across voxels.

### Association between visual featural effects and the domain effects in VOTC

To test whether the documented domain selectivity (e.g., animate vs. inanimate) might reflect visual featural effects, we examined if a voxel’s visual featural vector could predict its domain preference strength across different datasets. We adopted an independent dataset, where 31 healthy participants viewed blocks of gray-scale pictures of animals and artifacts, collapsing the two healthy control group data of published studies in our lab that used identical data acquisition procedures [35,36]. The data were preprocessed with the same procedure reported above and the preprocessed functional images of each participant were entered into a GLM, which included onset regressors for animate and inanimate together with 6 head motion regressors. Contrast images for animate vs. inanimate items were then calculated and averaged across participants to construct the animacy-domain selectivity VOTC map. In the linear regression model (in SPSS), the domain selectivity VOTC map (from this independent dataset) was treated as the dependent variable and the 17 full-model visual feature maps (from the main fMRI experiment) were included as independent variables. The adjusted R-squared of the regression model was reported.

To further test the effects of a tripartite structure [4], using the independent fMRI dataset, a GLM included onset regressors for animals, large artifacts, and small manipulable artifacts as well as 6 head motion regressors was constructed. Contrast images for each object domain with other two object domains were calculated and averaged across participants to construct three tripartite-domain selectivity VOTC maps. The associations between each of these maps and 17 full-model visual feature maps were then tested using the same procedure.

## Acknowledgments

We thank Shiguang Shan for discussions about computational vision models, and Joshua B. Julian for generously sharing codes for the right angle and curvature computation.

## Supporting Information

### S1 Text. Supplemental methods and results

**S1 Fig. Schematic of computational vision models for geometry/shape, Fourier power, and color-space visual features**. (A) Right angle and curvature. Step 1 shows the example of edge detection. Step 2 shows right-angle/curvature Gabor filters of one orientation in four spatial scales. Each Gabor filter had 16 orientations (22.5°-360° in 22.5° step). The curvature Gabor filter had 5 levels of curvature (π/256, π/128, π/64, π/32, π/16) and the level of curvature shown here is π/64. Step 3 shows example results of the convolution between edge maps and Gabor filters of different spatial scales. Step 4 shows the example max map of curvature/right angle across all orientations and levels of curvature in each spatial scale, which was then averaged across all pixels of each image and further z-scored across the image set. The resulting Gabor coefficients for each image were finally averaged across 4 spatial scales and z-scored to provide a single value for each image to represent the amount of curvature/right angle information in that image. (B) Fourier-power-space features. The Fourier transform of each image was first obtained. The mean energy over two spatial frequency bins (> 5 cycles/degree and < 1 cycles/degree) was calculated to represent the high and low spatial frequency energy of each image. The mean energy in each of 4 orientation bins (vertical (0°), left oblique (45°), horizontal (90°), and right oblique (135°); bandwidth: 20°) was calculated to represent energy in each orientation. (C) Color-space features. Pixel colors in each image were converted from RGB space into CIE-LCH space using the MATLAB “colorspace” package. The hue layer was divided into 8 bins with equal length, which started from the 338°-23° bin, in 45° steps, and roughly corresponded to red, orange, yellow, green, cyan, indigo, blue, and purple. The right panel shows the pixels in example image belonging to each bin of hues and pixel amplitudes in the luminance/chroma layer. In the hue layer, the numbers of pixels within object mask in each bin were counted as the hue-specific measures. The luminance and chroma of each image were calculated by averaging amplitudes within object mask in the luminance or chroma layer.

**S2 Fig. Descriptive statistics of the 20 visual feature weights across 95 object images**. (A)Distribution plots of z-transformed weights of each visual feature across 95 images. (B) Bar charts show mean and standard deviation of each visual feature weight broken down by tripartite domain structures (32 animate items, 28 large artifacts, 35 small manipulable artifacts). Red lines indicate significant difference between two bars. Blue asterisks indicate significant difference between animates (32 items) and inanimate artifacts (63 items).

**S3 Fig. The correlations among visual features**. The correlation matrix of 20 visual features obtained from computational vision models in 95 images. The color bar indicates the Pearson R value.

**S4 Fig. Independent-model parametric modulation results**. colors cyan and indigo, red and purple, were collapsed due to high correlation, resulting in 18 features. Each feature was entered into the parametric modulation model separately, yielding a beta value for each visual feature at each voxel. The activation maps are threshold at cluster-level FWE corrected *p* < .05 within the VOTC mask, with voxel-wise *p* < .001. The outlines show the domain-preferring clusters for large artifacts (bilateral PPA), animals (bilateral latFG), and small manipulable artifacts (left LOTC), localized by contrasting each object domain with the other two domains (see Materials and Methods).

**S5 Fig. Relationship between animacy-domain-model and visual featural topography**. The left column shows the “prototypical” visual feature vectors associated with animate domain or inanimate domain, respectively. Dots indicate that beta values were significant at FDR corrected *q* < .05 for 36 comparisons. The middle column shows the Pearson correlation maps between each of these “prototypical” vectors and VOTC voxels’ neural-feature vector obtained from the fMRI parametric modulation analyses. The correlation maps are thresholded at cluster-level FWE corrected *p* < .05, voxel-wise *p* < .001. Scatter plots show the correlations for the peak voxels. The right column presents the peak voxel’s neural-feature vectors.

**S6 Fig. Relationship between tripartite-domain-model and visual featural topography**. The left column shows the “prototypical” visual feature vectors associated with animals, large artifacts and small manipulable artifacts, respectively. Dots indicate that beta values were significant at FDR corrected *q* < .05 for 54 comparisons. The middle column shows the Pearson correlation maps between each of these “prototypical” vectors and VOTC voxels’ neural-feature vector obtained from the fMRI parametric modulation analyses. The correlation maps are thresholded at cluster-level FWE corrected *p* < .05, voxel-wise *p* < .001. Scatter plots show the correlations for the peak voxels. The right column presents the peak voxel’s neural-feature vectors.

**S7 Fig. Visual feature effects independent of object domains**. (A) Results of the visual featural effects with tripartite-domain structure regressed out. The same full-model parametric modulation procedure as in **Fig 2** was used, except that the visual feature weights of 95 images were residuals of the original visual features after regressing out the tripartite-domain structure. The bar plots show the averaged beta values of visual features across subjects in the ROIs that showed overlap between domain preferring regions (black line contours) and the featural maps in the original main analyses (see feature effects located in the domain-preferring outlines in **Fig 2**). (B) Results of visual featural effects in objects belonging to the non-preferring domains. In the parametric modulation analyses to obtain the visual featural effects, only object pictures belonging to the non-preferring domains were used. That is, for visual features significant in the large-artifact-preferring PPA, the parametric modulation model used pictures of animals and small manipulable artifacts. For visual features significant in animal-preferring latFG, the parametric modulation model used pictures of large and small artifacts. For visual features in small-artifact-preferring LOTC, the parametric modulation results used pictures of animals and large artifacts. The bar plots show the averaged beta values of visual features across subjects, in the same ROIs as in (A); error bars indicate standard error. Asterisks indicate that beta values were significantly greater than zero (one-tailed one-sample *t* tests, FDR corrected *q* < .05).

**S8 Fig. Visual featural effects without object context**. The brain activity maps for the contrasts between horizontal/vertical lines and curvature are thresholded at cluster-level FWE-corrected *p* < .05 within the VOTC mask, voxel-wise *p* < .001 and other contrast maps were thresholded at voxel-wise *p* < .05, cluster size > 270 mm^3^. The outlines show the domain-preferring clusters for animals (bilateral latFG), scenes (bilateral PPA), and small manipulable artifacts (left LOTC) localized by contrasting each domain with the other two domains in feature-validation fMRI experiment. Stimuli shapes are showing below the brain maps. Note in the actual experiment, each array contained more elements/dots than the examples shown here (See **S1 Text** for details).

**S9 Fig. The activation map in the VOTC by contrasting curved objects (e**.**g**., **button) with rectilinear objects (e**.**g**., **blackboard) in main fMRI experiment**. Previous studies have reported significant curvature effects in VOTC by contrasting round with rectilinear items (Yue et al., 2014). To replicate this result, we classified 95 object images in main fMRI experiment into 3 conditions: “curved items” (top 25% items sorted by curvature weights), “rectilinear items” (top 25% items sorted by right angle weights), and others. The nine items overlapped between “curved items” and “rectilinear items” (e.g., wrench) were recoded into the condition of other items, leaving 15 items in curved and rectilinear conditions. A GLM was then built including three conditions (“curved items”, “rectilinear items” and “other items”) and the contrast of “curved items” vs. “rectilinear items” was computed in individual subjects. Random-effects analysis was conducted using one-sample *t*-test (SnPM13). Thresholded at voxel-wise *p* < .01, the activation map shows that compared with rectilinear items, curved items activated the bilateral latFG, which overlapped with animal-preferring region. The outlines mean the same as in **S4 Fig**.

**S1 Table. Comparisons of visual feature weights across three object domains using ANOVA with post hoc contrasts between pairs of domains**.

**S2 Table. Logistic/Linear regression coefficients (p values) of visual feature weights for predicting domain model/relevance of response model in 767 images**.

## References

1. Felleman DJ, Van Essen DC. Distributed Hierarchical Processing in the Primate Cerebral Cortex. Cereb Cortex. 1991;1: 1–47. doi: 10.1093/cercor/1.1.1-a

2. Kriegeskorte N, Mur M, Ruff DA, Kiani R, Bodurka J, Esteky H, et al. Matching Categorical Object Representations in Inferior Temporal Cortex of Man and Monkey. Neuron. 2008;60: 1126–1141. doi: 10.1016/j.neuron.2008.10.043

3. Kanwisher N. Functional specificity in the human brain: A window into the functional architecture of the mind. PNAS. 2010;107: 11163–11170. doi: 10.1073/pnas.1005062107

4. Konkle T, Caramazza A. Tripartite Organization of the Ventral Stream by Animacy and Object Size. J Neurosci. 2013;33: 10235–10242. doi: 10.1523/JNEUROSCI.0983-13.2013

5. Grill-Spector K, Weiner KS. The functional architecture of the ventral temporal cortex and its role in categorization. Nat Rev Neurosci. 2014;15: 536–548. doi: 10.1038/nrn3747

6. Levy I, Hasson U, Avidan G, Hendler T, Malach R. Center–periphery organization of human object areas. Nat Neurosci. 2001;4: 533–539. doi: 10.1038/87490

7. Hasson U, Levy I, Behrmann M, Hendler T, Malach R. Eccentricity Bias as an Organizing Principle for Human High-Order Object Areas. Neuron. 2002;34: 479–490. doi: 10.1016/S0896-6273(02)00662-1

8. Bracci S, Beeck HO de. Dissociations and Associations between Shape and Category Representations in the Two Visual Pathways. J Neurosci. 2016;36: 432–444. doi: 10.1523/JNEUROSCI.2314-15.2016

9. Kaiser D, Azzalini DC, Peelen MV. Shape-independent object category responses revealed by MEG and fMRI decoding. Journal of Neurophysiology. 2016;115: 2246–2250. doi: 10.1152/jn.01074.2015

10. Proklova D, Kaiser D, Peelen MV. Disentangling Representations of Object Shape and Object Category in Human Visual Cortex: The Animate–Inanimate Distinction. Journal of Cognitive Neuroscience. 2016;28: 680–692. doi: 10.1162/jocn_a_00924

11. Bracci S, Ritchie JB, de Beeck HO. On the partnership between neural representations of object categories and visual features in the ventral visual pathway. Neuropsychologia. 2017;105: 153–164. doi: 10.1016/j.neuropsychologia.2017.06.010

12. Peelen MV, Downing PE. Category selectivity in human visual cortex: Beyond visual object recognition. Neuropsychologia. 2017 [cited 8 May 2017]. doi: 10.1016/j.neuropsychologia.2017.03.033

13. Long B, Yu C-P, Konkle T. Mid-level visual features underlie the high-level categorical organization of the ventral stream. PNAS. 2018; 201719616. doi: 10.1073/pnas.1719616115

14. Nasr S, Echavarria CE, Tootell RBH. Thinking Outside the Box: Rectilinear Shapes Selectively Activate Scene-Selective Cortex. J Neurosci. 2014;34: 6721–6735. doi: 10.1523/JNEUROSCI.4802-13.2014

15. Perrinet LU, Bednar JA. Edge co-occurrences can account for rapid categorization of natural versus animal images. Scientific Reports. 2015;5: 11400. doi: 10.1038/srep11400

16. Long B, Störmer VS, Alvarez GA. Mid-level perceptual features contain early cues to animacy. Journal of Vision. 2017;17: 20–20. doi: 10.1167/17.6.20

17. Yue X, Pourladian IS, Tootell RBH, Ungerleider LG. Curvature-processing network in macaque visual cortex. PNAS. 2014;111: E3467–E3475. doi: 10.1073/pnas.1412616111

18. Rosenthal I, Ratnasingam S, Haile T, Eastman S, Fuller-Deets J, Conway BR. Color statistics of objects, and color tuning of object cortex in macaque monkey. Journal of Vision. 2018;18: 1–1. doi: 10.1167/18.11.1

19. Chang L, Bao P, Tsao DY. The representation of colored objects in macaque color patches. Nature Communications. 2017;8: 2064. doi: 10.1038/s41467-017-01912-7

20. Bryan PB, Julian JB, Epstein RA. Rectilinear Edge Selectivity Is Insufficient to Explain the Category Selectivity of the Parahippocampal Place Area. Front Hum Neurosci. 2016;10. doi: 10.3389/fnhum.2016.00137

21. Mahon BZ, Caramazza A. What drives the organization of object knowledge in the brain? Trends in Cognitive Sciences. 2011;15: 97–103. doi: 10.1016/j.tics.2011.01.004

22. Abboud S, Maidenbaum S, Dehaene S, Amedi A. A number-form area in the blind. Nature Communications. 2015;6: 6026. doi: 10.1038/ncomms7026

23. Bi Y, Wang X, Caramazza A. Object Domain and Modality in the Ventral Visual Pathway. Trends in Cognitive Sciences. 2016;20: 282–290. doi: 10.1016/j.tics.2016.02.002

24. Osher DE, Saxe RR, Koldewyn K, Gabrieli JDE, Kanwisher N, Saygin ZM. Structural Connectivity Fingerprints Predict Cortical Selectivity for Multiple Visual Categories across Cortex. Cereb Cortex. 2016;26: 1668–1683. doi: 10.1093/cercor/bhu303

25. Brouwer GJ, Heeger DJ. Decoding and Reconstructing Color from Responses in Human Visual Cortex. J Neurosci. 2009;29: 13992–14003. doi: 10.1523/JNEUROSCI.3577-09.2009

26. Rajimehr R, Devaney KJ, Bilenko NY, Young JC, Tootell RBH. The “Parahippocampal Place Area” Responds Preferentially to High Spatial Frequencies in Humans and Monkeys. Whitney D, editor. PLoS Biology. 2011;9: e1000608. doi: 10.1371/journal.pbio.1000608

27. Troiani V, Stigliani A, Smith ME, Epstein RA. Multiple Object Properties Drive Scene-Selective Regions. Cereb Cortex. 2014;24: 883–897. doi: 10.1093/cercor/bhs364

28. Goffaux V, Duecker F, Hausfeld L, Schiltz C, Goebel R. Horizontal tuning for faces originates in high-level Fusiform Face Area. Neuropsychologia. 2016;81: 1–11. doi: 10.1016/j.neuropsychologia.2015.12.004

29. Zachariou V, Giacco ACD, Ungerleider LG, Yue X. Bottom-up processing of curvilinear visual features is sufficient for animate/inanimate object categorization. Journal of Vision. 2018;18: 3–3. doi: 10.1167/18.12.3

30. Hair JF, Black WC, Babin BJ, Anderson RE. Multivariate Data Analysis: Pearson New International Edition. Pearson Education Limited; 2013.

31. Wang X, Peelen MV, Han Z, He C, Caramazza A, Bi Y. How Visual Is the Visual Cortex? Comparing Connectional and Functional Fingerprints between Congenitally Blind and Sighted Individuals. J Neurosci. 2015;35: 12545–12559. doi: 10.1523/JNEUROSCI.3914-14.2015

32. Downing PE, Chan AW-Y, Peelen MV, Dodds CM, Kanwisher N. Domain Specificity in Visual Cortex. Cereb Cortex. 2006;16: 1453–1461. doi: 10.1093/cercor/bhj086

33. Moreno-Martínez FJ, Montoro PR. An Ecological Alternative to Snodgrass & Vanderwart: 360 High Quality Colour Images with Norms for Seven Psycholinguistic Variables. PLOS ONE. 2012;7: e37527. doi: 10.1371/journal.pone.0037527

34. Brodeur MB, Guérard K, Bouras M. Bank of Standardized Stimuli (BOSS) Phase II: 930 New Normative Photos. PLOS ONE. 2014;9: e106953. doi: 10.1371/journal.pone.0106953

35. He C, Peelen MV, Han Z, Lin N, Caramazza A, Bi Y. Selectivity for large nonmanipulable objects in scene-selective visual cortex does not require visual experience. NeuroImage. 2013;79: 1–9. doi: 10.1016/j.neuroimage.2013.04.051

36. Wang X, He C, Peelen MV, Zhong S, Gong G, Caramazza A, et al. Domain Selectivity in the Parahippocampal Gyrus Is Predicted by the Same Structural Connectivity Patterns in Blind and Sighted Individuals. J Neurosci. 2017;37: 4705–4716. doi: 10.1523/JNEUROSCI.3622-16.2017

37. Bao P, She L, McGill M, Tsao DY. A map of object space in primate inferotemporal cortex. Nature. 2020;583: 103–108. doi: 10.1038/s41586-020-2350-5

38. Nasr S, Tootell RBH. A Cardinal Orientation Bias in Scene-Selective Visual Cortex. J Neurosci. 2012;32: 14921–14926. doi: 10.1523/JNEUROSCI.2036-12.2012

39. Lescroart MD, Stansbury DE, Gallant JL. Fourier power, subjective distance, and object categories all provide plausible models of BOLD responses in scene-selective visual areas. Front Comput Neurosci. 2015;9. doi: 10.3389/fncom.2015.00135

40. Caldara R, Seghier ML, Rossion B, Lazeyras F, Michel C, Hauert C-A. The fusiform face area is tuned for curvilinear patterns with more high-contrasted elements in the upper part. NeuroImage. 2006;31: 313–319. doi: 10.1016/j.neuroimage.2005.12.011

41. Bracci S, Ritchie JB, Kalfas I, Beeck HPO de. The Ventral Visual Pathway Represents Animal Appearance over Animacy, Unlike Human Behavior and Deep Neural Networks. J Neurosci. 2019;39: 6513–6525. doi: 10.1523/JNEUROSCI.1714-18.2019

42. Sha L, Haxby JV, Abdi H, Guntupalli JS, Oosterhof NN, Halchenko YO, et al. The Animacy Continuum in the Human Ventral Vision Pathway. Journal of Cognitive Neuroscience. 2014;27: 665–678. doi: 10.1162/jocn_a_00733

43. Lafer-Sousa R, Conway BR, Kanwisher NG. Color-Biased Regions of the Ventral Visual Pathway Lie between Face- and Place-Selective Regions in Humans, as in Macaques. J Neurosci. 2016;36: 1682–1697. doi: 10.1523/JNEUROSCI.3164-15.2016

44. Canny J. A Computational Approach to Edge Detection. IEEE Transactions on Pattern Analysis and Machine Intelligence. 1986;PAMI-8: 679–698. doi: 10.1109/TPAMI.1986.4767851

45. Krüger N, Peters G, Malsburg CVD. Object Recognition with a Sparse and Autonomously Learned Representation Based on Banana Wavelets. Learned Representation Based on Banana Wavelets, Technical Report Ir-Ini 96-11 Institut Fur Neuroinformatik, Ruhr-Universitat Bochum. 1996.

46. Getreuer P. Colorspace Transformations. 2020 [cited 19 Mar 2020]. Available: https://www.mathworks.com/matlabcentral/fileexchange/28790

47. Yan C-G, Wang X-D, Zuo X-N, Zang Y-F. DPABI: Data Processing & Analysis for (Resting-State) Brain Imaging. Neuroinform. 2016;14: 339–351. doi: 10.1007/s12021-016-9299-4

