## Supplementary material for "Visual featural topography in the human ventral visual pathway": S1 Text

**S1 Text. Supplemental methods and results**

Previous studies suggested that specific featural effects persisted without object domain information (Nasr & Tootell, 2012; Yue et al., 2014). We validated whether the featural effects we observed were independent from the domain effects: 1) validation analyses of the main fMRI experiment, looking at featural effects while taking out the corresponding domain information; 2) a validation fMRI experiment in which the stimuli were arrays of lines and random dot patterns varying in hue, without any object contexts (see **S8 Fig** for sample stimulus shapes).

**Visual featural effects independent of object domain information in the main fMRI experiment**

**Visual featural effects regressing out the domain model.** We performed a ROI analysis computing the visual featural effects including the tripartite-domain model as covariates. The ROIs were defined as the overlapping voxels between the object-domain-preferring regions (bilateral latFG, bilateral PPA and left LOTC; see Materials and Method for details) and the significant clusters of each feature modulation in full-model parametric modulation (overlapping clusters containing at least 10 voxels were considered). We regressed out dummy-coded tripartite-domain model (assigning items from the corresponding domain as 1 and other items as 0) from each feature and re-ran the full model parametric modulation analysis using feature residuals as the modulating parameters. This procedure statistically eliminates the shared variance between feature weights and domains. The resulting parameter estimates for each feature in each ROI were averaged across voxels and compared with zero across participants using one-tailed one-sample *t*-tests, followed by FDR correction for multiple comparisons. The results (**S7A Fig**, FDR corrected *q* < .05) showed that in the 15 ROIs all effects remained similar to those before regressing out the tripartite-domain model.

**Visual featural effects in non-preferring domains.** We further examined the visual featural effects using object images belonging to the non-preferring domains to fully exclude any potential domain effects. For example, we tested whether the right-angle effect in PPA, the large-artifacts-preferring region, remained when participants were looking at images of animals and small manipulable artifacts. At the individual participant level, GLMs were built including two regressors coding stimuli as “preferring domain” items and “non-preferring domain” items, and for the “non-preferring domain” condition, visual feature weights were entered for the full-model parametric modulation analysis. As above, the resulting parameter estimates for each feature in the ROIs showing feature and domain overlap were averaged across voxels and compared with zero across participants using one-tailed one-sample *t*-tests, followed by FDR correction for multiple comparisons. The results (**S7B Fig**, FDR corrected *q* < .05) showed that effects in 10 of the 15 ROIs remained stable.

**Feature-validation fMRI experiment using isolated visual features**

**fMRI stimuli.** Stimuli consisted of 112 images from 14 conditions. Nine conditions were individual visual features: 4 different line-based features as shape arrays (8 arrays of right angles; 8 arrays of semicircles; 8 arrays of horizontal lines; 8 arrays of vertical lines), and the remaining 5 were hue-based features (40 random dot patterns in each of: red, orange, green, cyan, and blue). Another condition of grayscale random dot patterns was included as control for hue-based features. The remaining conditions (non-cardinal lines, phase-scrambled images of non-cardinal lines, and two types of combination of multiple visual features) were designed for other research purposes. Only the nine conditions of individual features and the grayscale random dot pattern condition were analyzed here. Each shape array contained 40 elements, and all the shape arrays were located randomly within a virtual circular limit (radius = 200 pixels) (constructed largely following a previous study (Nasr et al., 2014)). Orientations of shape arrays except for horizontal and vertical lines were varied semi-randomly to minimize orientation biases. Images of dot patterns were randomly distributed small dots (< 0.1°) rendered with gray or different color hues. For the colored dot patterns, we sampled 8 equally spaced points in (CIE) L*C*H space (Commission Internationale de l’Eclairage), that started from h = 0, at L* = 69, c* = 36, and for the following 5 hues: red (h = 0°), orange (h = 45°), green (h = 135°), cyan (h = 180°), blue (h = 270°). All stimuli had equal number of pixels and were presented against a Gaussian noise background (using the MATLAB function imnoise). The SHINE toolbox (Willenbockel et al., 2010) was used to match spatial frequency and luminance across all stimuli and to match the Fourier spectrums across dot conditions. The feature-validation fMRI experiment further included domain-functional localizer runs that included 90 black-and-white images taken from a previous study (He et al., 2013), including scenes (30 images), animals (30 images) and small manipulable objects (30 images). All images subtended 7.6° x 7.2° of visual angle (400 x 400 pixels).

**fMRI experiment design.** Participants viewed images of isolated shape arrays and dot patterns (described above). There were 7 runs in total, each lasting for 236 s. The stimuli were presented in 8-s blocks, separated by 8 s of fixation. Each block consisted of 8 images, each presented for 0.8 s and separated by 0.2 s of blank screen. Each condition appeared once in each run and the order of conditions was randomized. Each run started and ended with 10 s of fixation. Participants were instructed to pay attention to the stimuli and to press a button when a white dot (< 0.5°) appeared in the center of the screen, which occurred 0, 1 or 2 times per block with equal possibility. Six of the 19 participants also participated in a second scanning session of 9 runs on a separate day; we thus analyzed 16 runs for those 6 participants. After the isolated-feature runs, 2 domain localizer runs were conducted, in which participants viewed images from 3 domains (animals, scenes and small manipulable artifacts). Images were presented in 24 s blocks, separated by 8 s of fixation. Each block consisted of 30 images, each presented for 0.3 s and separated by 0.5 s blank screen. Each condition was repeated 3 times per run. During the localizer, participants performed a 1-back repetition detection task, pressing a button using their right index finger whenever two consecutive images were the same. There were 0, 1, or 2 times repetitions per block (same frequency for each condition).

**MRI acquisition.** Feature-validation fMRI experiment was conducted at the Imaging Center for MRI Research, Peking University, using a Siemens Prisma 3-T scanner with a 64-channel phase-array head coil. Functional data were collected with a simultaneous multi-slice (SMS) sequence (62 slices, multi-band factor = 2; TR = 2000 ms; TE = 30 ms; flip angle = 90°; matrix size = 112 𝗑 112; voxel size =2 𝗑 2 𝗑 2 mm with gap of 0.3 mm). T1-weighted anatomical images were acquired using a 3D MPRAGE sequence: 192 sagittal slices; 1 mm thickness; TR = 2530 ms; TE = 2.98 ms; inversion time = 1100 ms; flip angle = 7°; FOV = 256 × 224 mm; voxel size = 0.5 × 0.5 × 1 mm, interpolated; matrix size = 512 × 448.

**Data analysis.** Statistical analyses were carried out within a bilateral VOTC mask (Wang et al., 2015). All 14 stimulus conditions were included in the GLM as regressors, along with 6 head motion regressors. The following contrasts were examined: dot patterns in different hues vs. gray-scale dots, right angles vs. semicircles, horizontal lines vs. semicircles, and vertical lines vs. semicircles. For the domain localizer runs, three conditions (animals, scenes and small manipulable artifacts) were included in the GLM as regressors, along with 6 head motion regressors. Domain-preferring regions were identified by contrasting each domain with the other two. At the conventional threshold, the activation clusters of animals and scenes in this experiment spread to early visual cortex, so the localization of PPA and animal-latFG were constrained in an anatomical mask including bilateral temporal fusiform cortex, bilateral temporal occipital fusiform cortex and bilateral parahippocampal gyrus in the Harvard-Oxford atlas (<http://fsl.fmrib.ox.ac.uk/fsl/fslwiki/Atlases>).

**Results.** The results (**S8 Fig**) showed that semicircles, compared to horizontal/vertical lines or to right angles, activated bilateral or right latFG close to animal-preferring clusters. Right angles compared to semicircles activated bilateral medFG corresponding to PPA at a lenient threshold (voxel-wise p < .05, uncorrected, cluster size > 270 mm^3^). No effects of horizontal or vertical lines greater than semicircles were found in the fusiform gyrus. For dot patterns rendered with different hues (red, orange, green, cyan, and blue), compared to grayscale dot patterns, weak activations (voxel-wise p < .05, uncorrected, cluster size > 270 mm^3^) were observed posterior to the domain-preferring clusters or sandwiched between latFG and PPA. In summary, when presented as isolated single-feature stimuli without object contexts, the effects of right angles in PPA and curvature in right latFG tended to remain; the effects of right angles in LOTC and of hues and orientations broadly, disappeared.
