## Supplementary material for "Visual featural topography in the human ventral visual pathway": S1 Table

S1 Table. Comparisons of visual feature weights across three object domains using ANOVA with post hoc contrasts between pairs of domains.

|  |  |  |  | Tukey’s HSD Comparisons (*p* values) | | |
| --- | --- | --- | --- | --- | --- | --- |
| Features showing significant domain differences in ANOVA | F (*p*) | Domain | Mean (SD) | Large | Animal | Small |
| Right angle | 6.77 (0.002) | Large | 0.50 (1.10) |  | 0.001 | 0.065 |
|  |  | Animal | -0.39 (0.09) |  |  | 0.287 |
|  |  | Small | -0.04 (1.20) |  |  |  |
| Number of pixels | 16.37 (<0.001) | Large | 0.51 (0.92) |  | 0.530 | <0.001 |
|  |  | Animal | 0.27 (0.79) |  |  | <0.001 |
|  |  | Small | -0.66 (0.89) |  |  |  |
| Elongation | 15.47 (<0.001) | Large | -0.36 (0.39) |  | 0.984 | <0.001 |
|  |  | Animal | -0.40 (0.20) |  |  | <0.001 |
|  |  | Small | 0.65 (1.38) |  |  |  |
| Low spatial frequency | 6.59 (0.002) | Large | 0.32 (1.02) |  | 0.907 | 0.004 |
|  |  | Animal | 0.22 (0.82) |  |  | 0.012 |
|  |  | Small | -0.46 (0.99) |  |  |  |
| 0^o^ orientation | 6.21 (0.003) | Large | 0.47 (1.15) |  | 0.158 | 0.002 |
|  |  | Animal | 0.01 (0.76) |  |  | 0.214 |
|  |  | Small | -0.38 (0.93) |  |  |  |
| 90^o^ orientation | 5.08 (0.008) | Large | 0.35 (1.06) |  | 0.631 | 0.008 |
|  |  | Animal | 0.12 (0.71) |  |  | 0.076 |
|  |  | Small | -0.39 (1.07) |  |  |  |
| 135^o^ orientation | 8.06 (0.001) | Large | -0.09 (0.78) |  | 0.041 | 0.391 |
|  |  | Animal | 0.51 (1.30) |  |  | <0.001 |
|  |  | Small | -0.40 (0.56) |  |  |  |
| Orange | 5.11 (0.008) | Large | 0.03 (1.03) |  | 0.329 | 0.243 |
|  |  | Animal | 0.38 (1.14) |  |  | 0.005 |
|  |  | Small | -0.37 (0.68) |  |  |  |
| Yellow | 5.43 (0.006) | Large | 0.24 (1.45) |  | 0.997 | 0.021 |
|  |  | Animal | 0.26 (0.87) |  |  | 0.013 |
|  |  | Small | -0.42 (0.34) |  |  |  |

SD: Standard deviation
