## Supplementary material for "Visual featural topography in the human ventral visual pathway": S2 Table

S2 Table. Logistic/Linear regression coefficients (*p* values) of visual feature weights for predicting domain model/relevance of response model in 767 images.

|  |  | Dependent variable | | | |
| --- | --- | --- | --- | --- | --- |
| Independent variable | Animate | Inanimate | Relevance of Fight-or-Flight | Relevance of Navigation | Relevance of Manipulation |
| Right angle | -2.62 (<0.001) | 2.62 (<0.001) | -0.23 (<0.001) | 0.30 (<0.001) | -0.03 (0.401) |
| Curvature | -0.03 (0.635) | 0.03 (0.635) | -0.02 (0.677) | -0.06 (0.109) | 0.05 (0.143) |
| Number of pixels | 0.00 (0.969) | 0.00 (0.969) | -0.04 (0.292) | 0.34 (<0.001) | -0.33 (<0.001) |
| Elongation | -1.17 (<0.001) | 1.17 (<0.001) | -0.08 (0.023) | -0.17 (<0.001) | 0.27 (<0.001) |
| High spatial frequency | -0.01 (0.938) | 0.01 (0.938) | 0.04 (0.274) | 0.22 (<0.001) | -0.19 (<0.001) |
| Low spatial frequency | 0.20 (0.009) | -0.20 (0.009) | 0.07 (0.069) | 0.20 (<0.001) | -0.27 (<0.001) |
| 0^o^ orientation | -0.29 (<0.001) | 0.29 (<0.001) | -0.12 (0.001) | 0.39 (<0.001) | -0.24 (<0.001) |
| 45^o^ orientation | 0.39 (<0.001) | -0.39 (<0.001) | 0.12 (0.001) | -0.12 (0.001) | -0.03 (0.470) |
| 90^o^ orientation | 0.27 (<0.001) | -0.27 (<0.001) | 0.10 (0.004) | 0.19 (<0.001) | -0.30 (<0.001) |
| 135^o^ orientation | 0.20 (0.011) | -0.20 (0.011) | 0.07 (0.061) | -0.13 (<0.001) | 0.03 (0.341) |
| Red/Purple | 0.06 (0.458) | -0.06 (0.458) | 0.04 (0.306) | -0.01 (0.807) | -0.02 (0.639) |
| Orange | 0.09 (0.222) | -0.09 (0.222) | 0.01 (0.710) | 0.14 (<0.001) | -0.14 (<0.001) |
| Yellow | 0.38 (<0.001) | -0.38 (<0.001) | 0.18 (<0.001) | 0.10 (0.006) | -0.21 (<0.001) |
| Green | 0.06 (0.450) | -0.06 (0.450) | 0.04 (0.265) | -0.02 (0.539) | -0.01 (0.849) |
| Cyan/Indigo | 0.13 (0.122) | -0.13 (0.122) | 0.02 (0.536) | -0.01 (0.823) | -0.02 (0.497) |
| Blue | -0.10 (0.184) | 0.10 (0.184) | -0.09 (0.015) | 0.04 (0.224) | -0.01 (0.852) |
| Luminance | -0.12 (0.095) | 0.12 (0.095) | -0.09 (0.017) | 0.01 (0.770) | 0.05 (0.130) |
| Chroma | -0.07 (0.367) | 0.07 (0.367) | -0.04 (0.311) | -0.07 (0.046) | 0.12 (0.001) |
